# Unfamiliar conspecifics increase freezing and shift movement towards immobility in zebrafish, most clearly in males

**DOI:** 10.64898/2026.10.29.760420

**Authors:** David Lagman, Pavla Hubená, Svante Winberg, Madeleine Le Grevés

## Abstract

Familiar companions can buffer stress in zebrafish (*Danio rerio*), but whether an unfamiliar shoal is itself a stressor, and whether the sexes respond alike, is unclear. We exposed adult zebrafish to a familiar or an unfamiliar shoal for at least 75 min, video-tracked their behaviour over the last 60 min and then measured whole-body cortisol. In males, an unfamiliar shoal increased freezing (+14.5 percentage points), reduced swim bouts and shifted movement towards a frozen state (8% to 49% of frames). In females, the changes were smaller and not significant. With the sexes pooled, whole-body cortisol was 1.6-fold higher after exposure to the unfamiliar shoal. Sex differences were only found for swim bouts and movement states, not for freezing or cortisol. We conclude that an unfamiliar shoal is a likely stressor for zebrafish, with behavioural effects clearest in males. Thus, who a zebrafish shares its tank with is not a neutral background, and the familiarity of conspecifics should be considered whenever fish are placed in new social settings in the lab.

**Summary statement:** Exposure to an unfamiliar rather than a familiar shoal increases freezing in zebrafish, most clearly in males, and moderately raises cortisol, but cortisol and behaviour are not linked across individuals.

## Introduction

Shoaling protects fish from predators and supports mate choice (Brown et al., 2011), but this can also bring aggression due to hierarchy dynamics (Winberg and Hubená, 2026). For example, winning and losing fights with conspecifics shapes later social interactions (Dugatkin and Druen, 2004; Oliveira et al., 2011). Fish can also recognise familiar conspecifics (Griffiths, 2003) and kin (Olsén et al., 1998). Once established, hierarchies in a shoal save time and energy and reduce injury risk (Tibbetts et al., 2022). However, environmental disturbance can destabilise these hierarchies and, under some conditions, raise aggression (Sneddon et al., 2006). Familiarity therefore mediates shoaling decisions, reduces aggression in territorial species and affects mate choice (Ward and Hart, 2003). Notably, unfamiliar fish are also potential mates, so the balance of costs and benefits may differ between the sexes.

In zebrafish, conspecifics can dampen fear (Faustino et al., 2017) but can also transmit it (Akinrinade et al., 2023), and social buffering of stress occurs across fishes (Gilmour and Bard, 2022). Familiarity changes how zebrafish respond to a fearful companion (Silva et al., 2019). So far, most previous work has used conspecifics as a buffer against another stressor, such as alarm substance (Faustino et al., 2017), while whether an unfamiliar shoal can itself act as the stressor has received much less attention.

Zebrafish is now an established model for stress and anxiety research (Stewart et al., 2012; Kalueff et al., 2013). As in other teleosts, their cortisol response is controlled by the hypothalamic–pituitary– interrenal (HPI) axis but stress responses can differ between the sexes: females showed smaller stress responses than males in one study (Vera-Chang et al., 2019). Endogenous estradiol can shape sexually dimorphic anxiety through the HPI axis (Tao et al., 2025). However, behaviour does not always follow the hormones in animals: social stress raises anxiety-like behaviour equally in both sexes (Bozi et al., 2021), and sex may affect the form of the defensive response in rats (Gruene et al., 2015) and shoal preference in zebrafish (Velkey et al., 2022).

Here we asked whether (i) male and female zebrafish respond differently in behaviour to a familiar and an unfamiliar shoal and (ii) if the sexes differ in whole-body cortisol level after this experience. Adult zebrafish were exposed alone to a familiar or an unfamiliar shoal for at least 75 min while their behaviour was video tracked, after which whole-body cortisol was measured in the same fish.

## Materials and methods

### Animals and experimental design

Adult zebrafish of mixed sex (*Danio rerio* (Hamilton, 1822); AB strain; bred on 4 June 2024) were obtained from the zebrafish facility at Karolinska Institutet (Stockholm, Sweden). They were kept in group-housing tanks (3–9.5 L) on Aquaneering racks (San Marcos, USA) at ∼28 °C under a 14 h light:10 h dark cycle and were fed Zebrafeed (Sparos, Olhão, Portugal) and rotifers twice a day. Fish were 108–135 days old at the time of testing. Sex was identified visually before the experiments and confirmed by dissection afterwards. Focal fish were alone in the arena only during testing, three or four at a time in separate arenas (20 sessions between 20 September and 17 October 2024). Each focal fish was exposed to a Known or an Unknown stimulus shoal of four fish (two males and two females); six shoals were used, each under both treatments (Table S14). Known focal fish came from the same facility tank and rack as their shoal and were acclimatised to the test tank and room together with it. Unknown focal fish came from a separate rack system and were acclimatised apart from the shoal. Acclimatisation lasted 1 h on each of two days before testing. A transparent divider with holes let water and odour cues pass between focal fish and shoal. Fish were allocated to treatments as evenly as possible across testing days without formal randomisation (Table S14). Metrics were computed automatically and cortisol assays were not blinded. Behavioural experiments were run between 10:00 and 15:00 each day to limit diurnal variation in cortisol. Seventy fish were tested and two males (one per treatment) were excluded by tracking quality control (Table S6), giving Known/male n = 15, Known/female n = 18, Unknown/male n = 14 and Unknown/female n = 21 (n = 68 for behaviour; n = 65 for cortisol). The study was approved by the Uppsala Animal Ethics Committee (permit 5.218-13393/2023).

### Video tracking and behavioural metrics

Each focal compartment (13.9 cm × 16.2 cm) was filmed from above; the videos were cropped to the arena and resampled to 10 frames s□^1^ (ffmpeg) and trajectories were extracted with idtracker.ai (Romero-Ferrero et al., 2019). The final 60 min of tracking in each recording (36,000 frames; two recordings that continued after the test were cut at the last tracked frame) were analysed in six 10-min bins. Fish had been with the shoal for at least 15 min before the analysed window. Per fish and bin we computed freezing (% of time with smoothed velocity < 0.4 cm s□^1^ for ≥ 2 s), mean velocity, a social preference index (SPI = [time near − time far] / total time; near and far are the thirds of the arena width closest to and farthest from the shoal), entries into the near-shoal third, thigmotaxis (% of time within 1.5 cm of a wall), mean absolute turn angle and tortuosity. Swim and freeze bouts were counted per fish over the trial. The partitions between compartments were located for each session from the empty band in the tracked positions, and frames in which a fish’s track lay outside its own compartment (identity swaps with a neighbour) were removed. Calibration and quality control are in Supplementary Methods S1–S2.

### Whole-body cortisol

Immediately after the trial, fish were frozen and stored at −80 °C. Whole bodies were partially thawed on ice, weighed and homogenised in 1 mL ice-cold PBS (VWR VDI 12 homogeniser). Cortisol was extracted with ethyl acetate (5 mL; vortexed for 1 min and centrifuged at 2,000 g for 10 min; three to four rounds with the supernatants pooled), following Canavello et al. (2011) with minor adaptations. The pooled extract was dried under nitrogen at 40 °C, reconstituted in 1 mL PBS and held overnight at 4 °C. Whole-body cortisol was then measured in triplicate with a competitive cortisol ELISA kit (96 tests, catalogue no. 15642299; Fisher Scientific GTF AB) at a 1:5 sample dilution on three plates, with concentrations read from a four-parameter logistic standard curve (Boster ElisaInfo calculator), and expressed per g wet body weight.

### Statistics

The individual fish was the unit of replication. Behaviour over time was analysed with linear mixed-effects models in R 4.3.3 (lme4, lmerTest, emmeans) containing Treatment × Sex, a quadratic trend over time, and random intercepts for session and for individual fish. Within-sex effects were extracted with emmeans and p-values were corrected with the Benjamini–Hochberg (BH) procedure. Counts of swim and freezing bouts were analysed with Poisson mixed models with an observation-level random effect. Post hoc sensitivity analyses used glmmTMB with outcome-matched families (Supplementary Methods S3). Movement was also classified with a three-state Gaussian hidden Markov model (HMM; depmixS4) fitted to log-transformed smoothed velocity and absolute turn angle, and each fish’s occupancy of each state was analysed by beta regression. Cortisol was log□□-transformed and analysed with a censored (Tobit) regression (Treatment × Sex + body weight) in which the 11 samples that read below the lowest standard, two of them at the zero standard, were treated as left-censored at the detection limit (lowest standard × dilution ÷ body weight). Sensitivity checks, including arena position as a fixed effect, are in Tables S2, S11 and S12. Associations between cortisol and behaviour were tested with Spearman correlations and seven behavioural metrics were summarised by principal component analysis (PCA).

### Use of large language models

Large language models were used to help write the analysis code (R and Python) and to draft and edit the manuscript. The authors designed the study, reviewed and tested all code, checked all reported values and references against their sources, and take full responsibility for the content.

## Results and Discussion

### Unfamiliar shoals increase freezing, mainly in males

Males increased freezing in response to an unfamiliar shoal, whereas the increase in females was smaller and not significant. Males exposed to a Known shoal rarely froze (mean 1.1% of time) while males exposed to an Unknown shoal froze for 15.6% of time from the first 10 min onwards (Fig. 1A,E). The Unknown − Known difference in males was +14.5 percentage points (95% CI 6.3–22.6; P = 0.00090; BH-adjusted across five outcomes P = 0.0045). In females the difference was +4.8 points (P = 0.19) and the Treatment × Sex interaction was not significant (P = 0.082). Binomial refits and adding recording day or stimulus shoal as a random effect gave the same conclusion (male P ≤ 0.0022; Tables S10, S11), as did adding arena position as a fixed effect (+11.9 points; P = 0.0068; Table S11b). The male effect was significant at freezing thresholds of 0.3 cm s□^1^ and above but not at 0.2 cm s□^1^ (Table S5); 0.2 cm s□^1^ lies at the tracking noise floor, where a motionless fish rarely qualifies (Supplementary Methods S2). The behavioural change resembles the anxiety-like response to novelty in zebrafish that includes reduced exploration, more freezing and higher cortisol (Cachat et al., 2010). We therefore conclude here that a novel shoal can trigger a similar strong phenotype in male zebrafish. An unfamiliar shoal may provide stressful signals from both novel mating opportunities and potential conflicts over rank in the new social constellation. Social stress raises anxiety-like behaviour in both sexes (Bozi et al., 2021), which fits the weaker changes in the same direction that we saw in females. Dominant males are more aggressive than subordinates (Filby et al., 2010), and social rank and sex both affect brain monoamine activity in zebrafish (Dahlbom et al., 2012). Dominance increased reproductive success in males but not in females (Paull et al., 2010), so conflict over rank may carry higher costs for males and could explain their broader behavioural response seen here.

**Figure 1.**
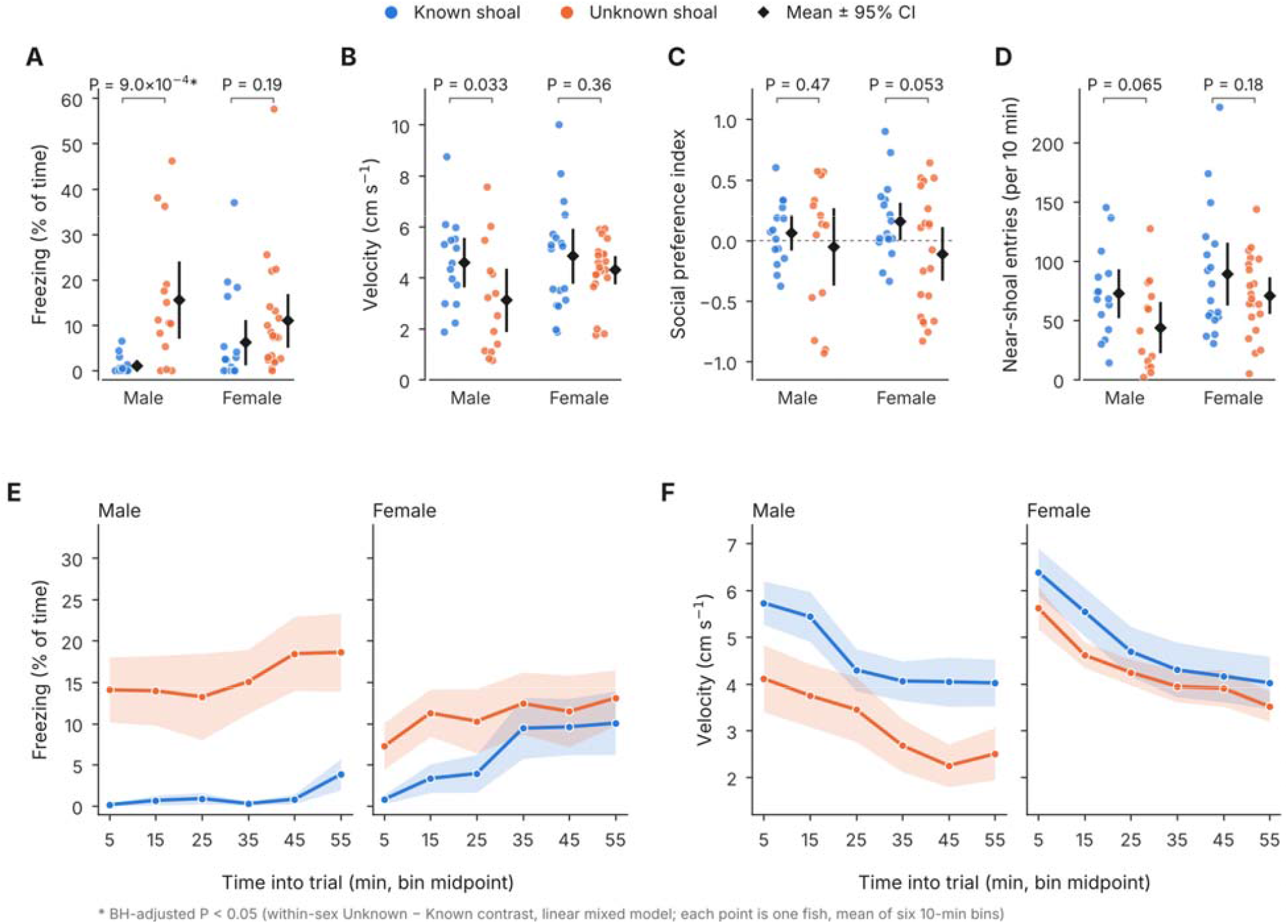
Unfamiliar shoals increase freezing in males; other behavioural measures show similar trends that are not robust. A–D) Per-fish means over the 60-min trial for freezing (A), velocity (B), social preference inde (SPI; positive = more time near the shoal; C) and entries into the near-shoal zone (D). Each point is one fish (Known shoal, blue; Unknown shoal, orange); diamonds and bars show mean ± 95% CI. P values are within-sex Unknown − Known contrasts from the linear mixed models; * BH-adjusted P < 0.05. Contrasts: freezing, males +14.5 percentage points, females +4.8; velocity, males −1.48 cm s□^1^, females −0.54; SPI, males −0.11, females −0.27; entries, males −28.8, females −18.2 (Table S1). E, F) Freezing (E) and velocity (F) across the trial; lines show mean ± s.e.m. of per-fish values in 10-min bins. Time courses of SPI and entries are in Fig. S3.

### Males shift their movement repertoire towards freezing

As shown above, Unknown-shoal males swam less and froze more. Their number of swim bouts was 0.35 times that of Known-shoal males (95% CI 0.24–0.52; P = 2.4 × 10□□), whereas freezing bouts increased with an imprecise ratio because Known-shoal males almost never froze (Table S3; Fig. S1). The swim-bout result was unchanged at peak-velocity cut-offs of 1 and 2 cm s□^1^ (Table S17). A threshold-free three-state HMM gave the same picture (Fig. 2). Mean occupancy of the frozen state rose from 8.0% to 48.7% of frames in males (beta regression; P = 9.0 × 10□□) and straight swimming fell from 65.4% to 33.2% (Tables S4, S13). The frozen state is a low-speed state rather than freezing in the strict sense: it also contains pauses shorter than 2 s and very slow movement (Supplementary Methods S3). In females the frozen state rose from 15.6% to 22.2% (P = 0.072) and the swim-bout reduction was smaller (Treatment × Sex ratio 2.55; P = 4.4 × 10□□). The straight-swim state also showed a significant Treatment × Sex interaction (P = 7.6 × 10□□) and the frozen state a borderline one (P = 0.059). Because these states are compositional (log-ratio test of the interaction P = 0.045; Table S4), the HMM gives the strongest but not unequivocal support for a sex difference.

**Figure 2.**
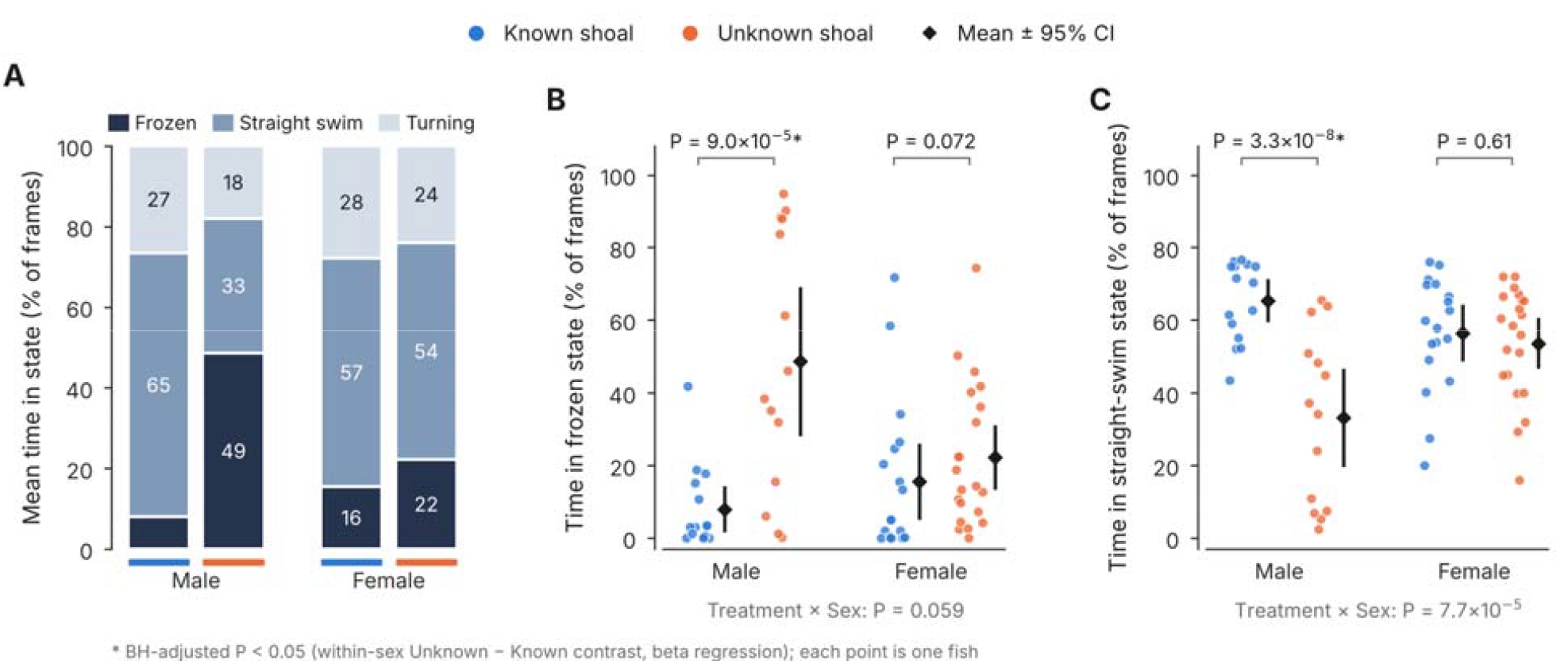
In males, an unfamiliar shoal shifts movement from straight swimming to a frozen state. A) Mean occupancy of three hidden Markov model states (frozen, straight swim, turning) by sex and treatment (blue bars, Known shoal; orange bars, Unknown shoal). B, C) Per-fish occupancy of the frozen (B) and straight-swim (C) states; each point is one fish, diamonds and bars show mean ± 95% CI. P values are within-sex Unknown − Known contrasts from beta regression (* BH-adjusted P < 0.05); interaction P values are for Treatment × Sex. Th classification does not rely on the freezing threshold used in Fig. 1.

### Velocity and social-zone measures change in the expected direction, but depend on the model

Although weak, measures of social-zone use changed in the direction expected if fish preferred the familiar shoal. In the linear models near-shoal entries were lower in Unknown-shoal fish (males −28.8, P = 0.065; females −18.2, P = 0.18) and velocity was lower in males (−1.48 cm s□^1^; P = 0.033), but neither survived BH correction (BH P = 0.11 and 0.082). SPI was lower after an Unknown shoal in both sexes, but not significantly (females −0.27, P = 0.053; males −0.11, P = 0.47). Thigmotaxis did not differ (Table S16). SPI also differed strongly between the left and right arenas (arena position P = 2.5 × 10□□; Table S11b), which were unevenly used across groups, so this measure should be read with care. Refitting entries and velocity with count and Gamma families (post hoc, because residuals were heavy-tailed) gave male reductions that survived BH correction (entries 0.44 times, P = 0.0058; velocity 0.56 times, P = 0.0021; Table S10), so we read these as model-dependent. The Treatment × Sex interaction was significant for velocity in the Gamma model (P = 0.039) and on a log scale (P = 0.041; Table S11), but not for entries (P ≥ 0.14). Zebrafish prefer familiar or established shoals at close range (Swaney et al., 2025; Velkey et al., 2022), and loss of buffering by familiar companions may contribute (Faustino et al., 2017; Gilmour and Bard, 2022). Sex-specific responses to unfamiliar fish occur in other species: mated female guppies prefer unfamiliar, unrelated males (Daniel and Rodd, 2016) and female rainbowfish avoid male relatives (Arnold, 2000). Adult female zebrafish prefer the odour of unrelated males whereas males show no preference by relatedness (Gerlach and Lysiak, 2006). Our SPI data do not separate the sexes, so whether zebrafish show such a pattern remains open.

### Whole-body cortisol is higher after an unfamiliar shoal

Whole-body cortisol was about 1.6-fold higher after exposure to an Unknown shoal (95% CI 1.05– 2.4; P = 0.030). In males the increase was 1.9-fold (95% CI 1.0–3.6; P = 0.045; Fig. 3A) and in females 1.3-fold (95% CI 0.76–2.3; P = 0.31), and the Treatment × Sex interaction was not significant (P = 0.40). These estimates treat the 11 samples that read below the lowest ELISA standard as left-censored at the detection limit. Eight of them are Known-shoal males, including K18 and K19, which read at the zero standard (extrapolated 5 and 19 pg g□^1^, more than ten times below any other fish). In males, censoring only K18 and K19 gave a 2.1-fold increase (P = 0.019), keeping all extrapolated values 3.6-fold (P = 0.006; Table S2) and excluding K18 and K19 1.85-fold (P = 0.078; Table S12), so the significance of this moderate effect depends on how values below the assay range are handled. Baseline cortisol tended to be higher in females (median 1,602 versus 871 pg g□^1^ after a Known shoal, the male median being below the calibrated range; P = 0.56), so a female ceiling cannot be excluded. Male-biased increases in cortisol have been reported after chronic stress in zebrafish (Rambo et al., 2017). Estradiol acts on anxiety through the HPI axis (Tao et al., 2025), and our design cannot separate such effects from sex differences in baseline.

**Figure 3.**
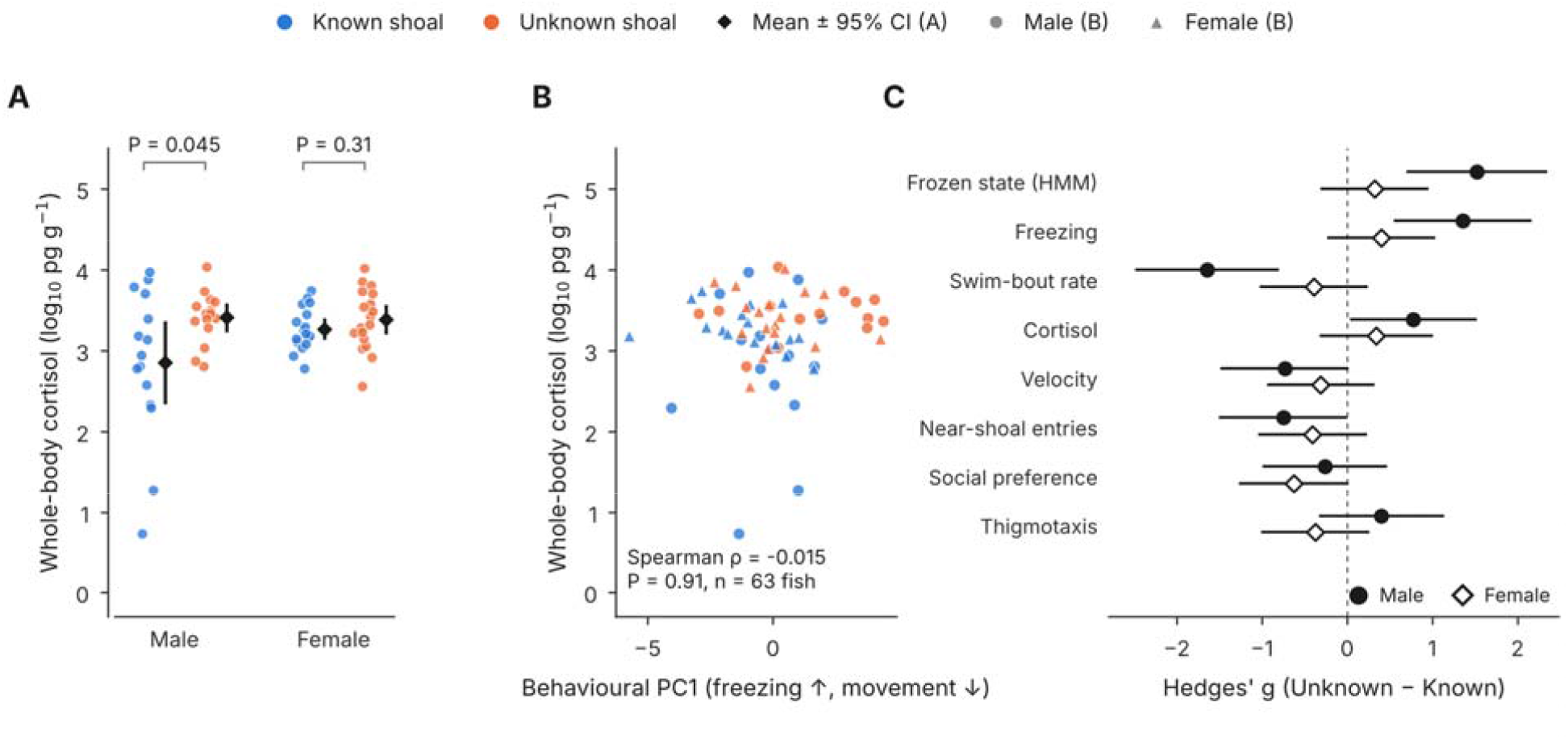
Cortisol is higher after an unfamiliar shoal, is unrelated to behaviour across individuals, and effects across outcomes are largest in males. A) log□□ whole-body cortisol (pg g□^1^ wet body weight) after at least 75 min of exposure to a Known (blue) or Unknown (orange) shoal; each point is one fish, diamonds and bars show mean ± 95% CI. Unknown − Known from a censored regression: males ≈ 1.9-fold (P = 0.045), females ≈ 1.3-fold (n.s.); Treatment × Sex P = 0.40 (n = 65), with the 11 samples below the lowest standard censored. Those samples are shown at their extrapolated values; the two lowest points (Known-shoal males) read at the zero standard. B) Cortisol against behavioural PC1 (males, circles; females, triangles; Spearman ρ= −0.015, P = 0.91; n = 63). C) Standardised effect sizes (Hedges’ g of Unknown − Known, with 95% CI) calculated descriptively from per-fish values for each outcome, by sex; positive values indicate higher values under the Unknown shoal.

### Cortisol and behaviour are not associated within individuals

Cortisol was uncorrelated with freezing, SPI, entries, velocity, thigmotaxis and PC1 of a PCA of behaviour (all |Spearman ρ| ≤ 0.13; all P ≥ 0.32; n = 63; Fig. 3B; Table S8; Fig. S2), also after removing the effects of treatment and sex (partial |ρ| ≤ 0.15; Table S12). Standardised effects across outcomes (Fig. 3C) were largest for freezing, frozen-state occupancy and swim-bout rate in males. Locomotor and immobility measures do not always track other indicators of anxiety (Johnson et al., 2023), and whole-body cortisol depends on social housing and was only weakly related to anxiety-test behaviour in an earlier study (Onarheim et al., 2022).

### Scope and future directions

With 14–21 fish per group the design detected the large male freezing effect, and larger samples should refine the sex comparison (Table S15). Cortisol was by design measured once, at the end of exposure, and focal fish were alone only during testing, whereas social isolation itself can alter cortisol (Shams et al., 2017). Because the divider passed odour as well as sight, future work can separate the sensory channels.

## Conclusions

Zebrafish exposed to an unfamiliar rather than a familiar shoal freeze more and shift their movement towards a frozen state, a robust effect most clearly seen in males. Whole-body cortisol was about 1.6-fold higher after exposure to the unfamiliar shoal. To our knowledge, this is the first study to combine cortisol and a broad behavioural panel in zebrafish of both sexes under familiar and unfamiliar conditions. The results show that social novelty alone, without a predator or alarm cue, can evoke a stress-like response, complementing the buffering role of familiar companions. Cortisol and behaviour were unrelated across individuals and capture different aspects of the response, so stress assays benefit from combining them and from including both sexes. Thus, who a zebrafish shares its tank with is not a neutral background: the familiarity of conspecifics should be considered whenever fish are moved, regrouped or tested in new social settings, in husbandry as much as in behavioural and stress research.

## Acknowledgements

We thank the Uppsala University Behavioural Facility (UUBF) for providing the behavioural equipment.

## Data and resource availability

Trajectory data, per-fish cortisol data, analysis code (R and Python) and all supplementary output tables are available at figshare: 10.6084/m9.figshare.34814397. No new biological materials were generated.

