## Supplementary text, figs and tables for "Unfamiliar conspecifics increase freezing and shift movement towards immobility in zebrafish, most clearly in males"

**Supplementary Methods**

***S1. Tracking, calibration, arena geometry and quality control***

Arenas were filmed from above, the videos were cropped to the arena and resampled to 10 frames s⁻¹ (ffmpeg; ConvertVideoFiles.sh), and per-frame x,y coordinates were extracted with idtracker.ai. Coordinates were converted from pixels to cm using the arena height (13.9 cm = 180 px; 0.0772 cm px⁻¹). The final 36,000 tracked frames (60 min) of each session were analysed in six 10-min bins (bins 0–5). Arena extent was estimated for each session from the pooled positions of the three or four fish (data-driven geometry), so that small between-session shifts of the camera were accommodated. The partitions between compartments were located as the empty band in the pooled positions (the partition plus the fish's half-width; 1.6–2.9 cm wide in every session); its edges are the inner walls of the neighbouring compartments (Table S7). Frames in which a fish's track lay outside its own compartment (identity swaps with a neighbour) were removed before any metric was computed (median 1.4% of frames per fish; up to 26%). Fish identities were checked for swaps; fish with more than 60 frames flagged as identity swaps, or any single-frame step above 38.6 cm (500 px), were excluded (the second criterion cannot occur within one compartment). Two of 70 fish were excluded (Table S6), leaving 68 fish. Tracking covered 36,271–37,939 frames per recording (60.5–63.2 min). Two recordings (Trials 5 and 14) continued filming for 6.3 and 8.4 min after the test; these untracked frames were removed before the final 36,000 frames were taken, so the first 0.5–3.2 min of tracking were not used. Six of 408 fish-bins (six fish) had fewer than 3,000 tracked frames (< 50% of the bin); all bins were retained.

***S2. Behavioural metrics***

Per fish and 10-min bin: freezing (% of frames in bouts of smoothed velocity < 0.4 cm s⁻¹ lasting ≥ 20 frames, i.e. ≥ 2 s; velocity smoothed with a 5-frame rolling window); mean velocity (cm s⁻¹); SPI = (time near − time far) / total time; entries into the near-shoal third of the arena; thigmotaxis (% of frames within 1.5 cm of a wall of the fish's own arena quadrant, including the inter-quadrant partition and the wall facing the stimulus shoal; a variant excluding the shoal-facing wall is analysed in Table S16); mean absolute turn angle (turns defined only for steps above a minimum displacement, otherwise set to 0); tortuosity (path length / net displacement). Per fish over the trial: number and duration of swim bouts (peaks in the smoothed velocity trace; pracma::findpeaks) and freezing bouts, and latency to enter the near-shoal zone. Tracking noise: the positional noise of a motionless fish, estimated from each fish's stillest 5-s windows (column noise_sd_px_still), was 0.13 px (median in both Unknown-shoal groups), an apparent speed of about 0.18 cm s⁻¹. The 0.4 cm s⁻¹ freezing threshold is therefore above the noise floor, whereas 0.2 cm s⁻¹ is at it (Table S5). Known-shoal males gave higher values (median 0.33 px) because their stillest windows still contained movement.

***S3. Statistical models***

Behavioural time series (lmerTest::lmer, REML): outcome ~ Treatment × Sex + poly(bin, 2) + (1|Session) + (1|fish_id), where fish_id is a unique identifier for each animal (session × individual index), so that repeated bins are nested within fish, the unit of replication. Random-effect terms with variance below 10⁻⁴ × residual variance were dropped and the model refitted; the session term was dropped for freezing, entries, SPI and velocity and retained for thigmotaxis, and the fish term was always retained. Dropping the empty session term left fixed-effect estimates, standard errors, degrees of freedom and p-values unchanged. p-values used Satterthwaite degrees of freedom. BH correction was applied within each effect family (Treatment, Sex, Treatment × Sex) across the five outcomes and across all 15 tests. Within-sex contrasts (Unknown − Known) were extracted with emmeans and BH-corrected across the ten contrasts. Treatment is referenced to the Known shoal and Sex to males, so the Treatment coefficient is the male Unknown − Known effect and Treatment × Sex is the additional female-specific difference. Bout counts per fish were analysed with Poisson mixed models with an offset for observation time and an observation-level random effect. HMM: a three-state Gaussian hidden Markov model (depmixS4) fitted to all frames of log1p(smoothed velocity) and absolute turn angle; states were labelled from their emission parameters as frozen, straight swim and turning. The frozen state is a low-speed state: its frames had a mean smoothed speed of 0.38 cm s⁻¹ (90th percentile 0.69 cm s⁻¹), and 38% of them met the 2-s freezing rule; the rest were shorter pauses and very slow movement. Three random starts and an independent implementation (hmmlearn) gave the same states (per-fish occupancy within 0.0003). For each state, each fish's occupancy proportion was analysed by beta regression (proportion ~ Treatment × Sex, session random intercept where estimable), BH-corrected across the three states, with within-sex contrasts and a Wilcoxon check; frame-level counts were not modelled because frames are autocorrelated. Cortisol: log₁₀(cortisol per g) ~ Treatment × Sex + body weight. The primary model is a left-censored (Tobit) regression (survival::survreg, Gaussian errors) in which the 11 samples that read below the lowest standard (50 pg mL⁻¹; K12, K14, K18, K19, K21, K25, K28, K31, OK7, OK16, OK18) are censored at the detection limit, 250 pg ÷ body weight (Tables S2b, S12); the same model fitted by ordinary least squares to the extrapolated values, with Type-III ANOVA (car::Anova), is given in Table S2. PCA: z-scored per-fish means of seven metrics (freezing, near-shoal entries, SPI, thigmotaxis, velocity, mean absolute turn, log₁₀ tortuosity), PC1 sign fixed so that the freezing loading is positive. Sensitivity refits (glmmTMB, same fixed effects and random effects (1|Session) + (1|fish_id)): Gaussian (REML); binomial, binomial with an observation-level random effect, or beta-binomial for freezing and thigmotaxis with frames per bin as trials; Poisson with observation-level random effect, or negative binomial, for entries (offset log frames); beta for SPI rescaled to 0–1; Gamma with log link for velocity. Coefficients are on the link scale and Wald (asymptotic) tests were used. Arena position (quadrant) was added as a fixed effect to the behavioural and HMM models (Table S11b). Cortisol sensitivity: HC3 robust errors, a permutation test of the male contrast (20,000 permutations), Wilcoxon tests, and refits excluding the lowest one or two values. Cortisol–behaviour associations were also tested as Spearman correlations between residuals after removing Treatment × Sex. Associations between cortisol and behaviour used Spearman and Pearson correlations. The original v3 run used R 4.5.0 with lme4 1.1-37, lmerTest 3.1-3, glmmTMB 1.1.14, emmeans 2.0.3, depmixS4 1.5-1. All numbers in this version (pipeline v3.4) were produced in R 4.3.3 with lme4 1.1-35.1, lmerTest 3.1-3, glmmTMB 1.1.8, emmeans 1.10.0, survival 3.5-8 and depmixS4 1.5-1; the unmodified v3 pipeline reproduced its R 4.5.0 outputs exactly in that environment.

**Supplementary Results**

***Table S1. Behavioural linear mixed models (code output: TableS01a, S01b, S01c)***

Unknown − Known estimates (95% CI where shown). Within-sex BH is across ten contrasts; interaction BH is within the Treatment × Sex family across five outcomes. Freezing and thigmotaxis in % of time, velocity in cm s⁻¹, entries as counts per 10-min bin.

| **Outcome** | **Males: estimate [95% CI]** | **P (BH)** | **Females: estimate** | **P (BH)** | **Treatment × Sex** | **P (BH)** |
| --- | --- | --- | --- | --- | --- | --- |
| Freezing | +14.47 [6.33, 22.60] | 0.00090 (0.0090) | +4.78 | 0.19 (0.31) | −9.69 | 0.082 (0.28) |
| Velocity | −1.48 [−2.80, −0.15] | 0.033 (0.16) | −0.54 | 0.36 (0.40) | +0.93 | 0.30 (0.50) |
| Entries near shoal | −28.84 [−58.92, 1.25] | 0.065 (0.16) | −18.18 | 0.18 (0.31) | +10.65 | 0.60 (0.60) |
| SPI | −0.114 [−0.422, 0.194] | 0.47 (0.47) | −0.268 | 0.053 (0.16) | −0.154 | 0.46 (0.58) |
| Thigmotaxis | +6.22 [−4.63, 17.07] | 0.28 (0.35) | −5.64 | 0.25 (0.35) | −11.86 | 0.11 (0.28) |

The linear trend over time was significant for all five outcomes (P ≤ 3.5 × 10⁻³). Group means per fish (TableS01d_descriptives): freezing 1.1 / 6.2 / 15.6 / 11.0% and velocity 4.60 / 4.85 / 3.12 / 4.31 cm s⁻¹ (Known male / Known female / Unknown male / Unknown female); SPI 0.06 / 0.16 / −0.05 / −0.11; entries 72.6 / 89.1 / 43.8 / 70.9; thigmotaxis 38.3 / 47.3 / 44.5 / 41.6%.

BH-adjusted P for the Treatment effect across the five outcomes (TableS01c_FDR_BH): freezing 0.0045, velocity 0.082, entries 0.11, SPI 0.47, thigmotaxis 0.33; across all 15 tests, freezing 0.013.

***Table S2. Cortisol models, n = 65 (code output: TableS02a, S02b, S02c)***

| **Term** | **Estimate (log₁₀)** | **95% CI** | **P** |
| --- | --- | --- | --- |
| Treatment (Unknown − Known), males | 0.553 (≈ 3.6-fold) | 0.164, 0.942 | 0.006 |
| Sex (female − male), Known shoal | 0.362 | −0.139, 0.863 | 0.153 |
| Treatment × Sex | −0.434 | −0.965, 0.097 | 0.107 |
| Body weight (g) | 0.235 | −1.214, 1.685 | 0.746 |
| Type-III ANOVA: Treatment | F₁,₆₀ = 6.44 | — | 0.014 |
| Type-III ANOVA: Sex | F₁,₆₀ = 0.51 | — | 0.476 |
| Type-III ANOVA: Treatment × Sex | F₁,₆₀ = 2.68 | — | 0.107 |

Cortisol per g (pg g⁻¹), medians (n): Known male 871 (15), Known female 1602 (17), Unknown male 2859 (15), Unknown female 2373 (18). Mean log₁₀: 2.85, 3.27, 3.41, 3.38. Eleven samples read below the lowest standard, eight of them in the Known-male group (two at the zero standard; Fig. 3A). This table keeps their extrapolated values; the primary analysis, with those samples censored, is in Table S2b.

***Table S2b. Primary cortisol model, n = 65 (code output: TableS02e_cortisol_tobit_primary)***

Left-censored (Tobit) regression of log₁₀ cortisol per g on Treatment × Sex + body weight; the 11 samples below the lowest standard are censored at 250 pg ÷ body weight. Fold changes are 10 to the power of the log₁₀ difference.

| **Contrast** | **Fold change [95% CI]** | **log₁₀ difference (SE)** | **P** |
| --- | --- | --- | --- |
| Unknown − Known, averaged over sexes | 1.60 [1.05, 2.45] | 0.205 (0.094) | 0.030 |
| Unknown − Known, males | 1.92 [1.01, 3.63] | 0.283 (0.141) | 0.045 |
| Unknown − Known, females | 1.34 [0.76, 2.35] | 0.126 (0.125) | 0.31 |
| Treatment × Sex (female minus male effect) | 0.70 [0.30, 1.63] | −0.158 (0.189) | 0.40 |
| Female − male, Known shoal | 1.27 [0.57, 2.84] | 0.103 (0.179) | 0.56 |

***Table S3. Bout-rate models (code output: TableS03_bout_rates)***

| **Outcome** | **Term** | **IRR [95% CI]** | **P** |
| --- | --- | --- | --- |
| Swim bouts | Unknown vs Known (males) | 0.35 [0.24, 0.52] | 2.4 × 10⁻⁷ |
| Swim bouts | Treatment × Sex | 2.55 [1.51, 4.30] | 4.4 × 10⁻⁴ |
| Freezing bouts | Unknown vs Known (males) | 36.3 [7.2, 183.3] | 1.4 × 10⁻⁵ |
| Freezing bouts | Treatment × Sex | 0.149 [0.018, 1.236] | 0.078 |

The wide interval for freezing bouts reflects near-zero freezing in Known-shoal males, which makes the ratio unstable; it should be read qualitatively. Using total or active observation time as offset gave swim-bout IRR 0.35 and 0.42 (interaction 2.55 and 2.26; Table S5).

***Table S4. HMM state occupancy (code output: TableS04a–e)***

| **State** | **Known M** | **Known F** | **Unknown M** | **Unknown F** | **Males: P (BH)** | **Females: P (BH)** | **Treatment × Sex P (BH)** |
| --- | --- | --- | --- | --- | --- | --- | --- |
| Frozen | 8.0% | 15.6% | 48.7% | 22.2% | 9.0×10⁻⁵ (2.7×10⁻⁴) | 0.072 (0.11) | 0.059 (0.059) |
| Straight swim | 65.4% | 56.5% | 33.2% | 53.7% | 3.3×10⁻⁸ (2.0×10⁻⁷) | 0.61 (0.61) | 7.6×10⁻⁵ (2.3×10⁻⁴) |
| Turning | 26.6% | 27.9% | 18.1% | 24.1% | 4.2×10⁻⁴ (8.3×10⁻⁴) | 0.31 (0.37) | 0.035 (0.052) |

Mean per-fish occupancy; beta regression on logit scale; contrasts are Unknown − Known. The Wilcoxon check agreed (males: frozen P = 0.002, straight P = 0.0004, turning P = 0.058; females: all P ≥ 0.10). Because the states are compositional, an isometric log-ratio analysis gave Treatment (males) P = 4.5 × 10⁻⁵ and Treatment × Sex P = 0.045 for the first coordinate, and Treatment P = 8.1 × 10⁻⁴ and Treatment × Sex P = 0.21 for the second.

***Table S5. Sensitivity analyses (code output: TableS05a–d)***

| **Analysis** | **Specification** | **Result** |
| --- | --- | --- |
| Freezing threshold | 0.2 / 0.3 / 0.4 (primary) / 0.6 / 0.8 cm s⁻¹ | Male effect +0.4 (P = 0.083) / +4.4 (P = 0.024) / +14.5 (P = 0.00090) / +31.4 (P = 3.1×10⁻⁵) / +37.7 (P = 1.4×10⁻⁵); female effect not significant at any threshold (P ≥ 0.12) |
| Random slopes | Random slope for time added (male Unknown − Known P from emmeans with its default degrees of freedom; random intercept → random slope) | Freezing 0.0012 → 8.6×10⁻⁴; velocity 0.037 → 0.040; entries 0.072 → 0.035; SPI 0.48 → 0.61; thigmotaxis 0.28 → 0.37 |
| Stimulus shoal | Add (1\|Shoal) to freezing and bout models | Freezing Treatment +14.5, P = 0.00090 (unchanged); swim-bout interaction IRR 2.55, P = 4.4×10⁻⁴ (unchanged); freezing-bout interaction IRR 0.149, P = 0.078 (unchanged) |
| Cortisol: random effects | Session; session + extraction batch; session + batch + shoal | Treatment effect 0.561 (P = 0.0051); 0.561 (P = 0.0049); 0.511 (P = 0.011) |
| Cortisol: weight adjustment | log₁₀(cortisol) ~ Treatment × Sex + log₁₀(weight) | Treatment 0.553, P = 0.0061 |
| Bout offsets | Total vs active time as offset | Swim-bout IRR 0.35 (P = 2.4×10⁻⁷) vs 0.42 (P = 9.2×10⁻⁸) |

Stimulus shoal identity was coded case-insensitively (Stim2 = stim2, Stim4 = stim4, Stim5 = stim5; confirmed by the authors).

***Table S6. Fish excluded by tracking quality control (code output: TableS06_excluded_fish)***

| **Fish** | **Session** | **Individual** | **Treatment** | **Sex** | **Stimulus shoal** | **Arena** | **Swap-flagged frames** | **Maximum step (cm)** |
| --- | --- | --- | --- | --- | --- | --- | --- | --- |
| K8 | Trial_4 | 1 | Known | Male | stim4 | left, bottom | 62 | 27.7 |
| OK7 | Trial_4 | 3 | Unknown | Male | stim1 | left, top | 76 | 29.6 |

Both fish were tested together on 26 September 2024 (Trial 4) and were excluded because more than 60 frames had an implausibly large single-frame displacement (> 3.86 cm; a possible identity swap), the exclusion threshold. Their identities were assigned from the median position of each tracked individual in the arena (individual 1, lower left; individual 3, upper left). The other two fish in the session (K7 and OK8) were retained.

***Table S7. Per-session arena geometry in cm (code output: TableS07_session_geometry)***

| **Session** | **x min** | **x midline** | **x max** | **y min** | **y midline** | **y max** |
| --- | --- | --- | --- | --- | --- | --- |
| Trial_1 | 1.74 | 16.34 | 30.19 | 2.14 | 14.09 | 26.82 |
| Trial_2 | 2.15 | 16.30 | 30.06 | 3.71 | 14.06 | 26.07 |
| Trial_3 | 1.57 | 15.77 | 30.21 | 2.29 | 14.29 | 26.29 |
| Trial_4 | 1.42 | 15.82 | 29.99 | 2.82 | 14.22 | 26.85 |
| Trial_5 | 1.62 | 15.92 | 30.32 | 2.21 | 14.30 | 26.77 |
| Trial_6 | 2.04 | 15.84 | 29.78 | 3.19 | 14.29 | 27.03 |
| Trial_7 | 1.93 | 16.28 | 30.45 | 2.30 | 14.45 | 26.69 |
| Trial_8 | 1.84 | 15.79 | 30.19 | 4.66 | 14.56 | 26.51 |
| Trial_9 | 1.87 | 15.81 | 30.38 | 2.67 | 14.42 | 27.20 |
| Trial_10 | 1.63 | 15.83 | 29.99 | 3.72 | 14.57 | 26.91 |
| Trial_11 | 1.43 | 16.18 | 30.56 | 2.46 | 14.36 | 27.06 |
| Trial_12 | 1.78 | 15.98 | 30.79 | 1.78 | 14.48 | 27.34 |
| Trial_13 | 1.56 | 16.16 | 30.54 | 1.40 | 14.35 | 26.94 |
| Trial_14 | 1.48 | 16.08 | 30.41 | 1.77 | 13.97 | 26.78 |
| Trial_15 | 1.88 | 16.08 | 30.61 | 2.56 | 14.21 | 26.92 |
| Trial_16 | 1.39 | 16.29 | 30.58 | 1.73 | 14.33 | 27.24 |
| Trial_17 | 1.42 | 15.92 | 29.29 | 3.33 | 14.78 | 27.12 |
| Trial_18 | 1.57 | 15.87 | 29.74 | 2.94 | 15.04 | 27.04 |
| Trial_19 | 1.56 | 15.86 | 29.42 | 3.43 | 15.03 | 27.27 |
| Trial_20 | 1.59 | 15.79 | 29.50 | 3.73 | 15.28 | 27.32 |

Drift of each boundary from its nominal position is given in the code output.

***Table S8. Cortisol–behaviour correlations, n = 63 (code output: TableS08_cortisol_behaviour)***

| **Metric** | **Spearman ρ** | **P** | **Pearson r** | **P** |
| --- | --- | --- | --- | --- |
| Freezing | 0.126 | 0.323 | 0.132 | 0.30 |
| SPI | −0.107 | 0.405 | −0.066 | 0.61 |
| Entries near shoal | −0.002 | 0.988 | 0.023 | 0.86 |
| Velocity | 0.031 | 0.811 | −0.067 | 0.60 |
| Thigmotaxis | 0.050 | 0.697 | 0.026 | 0.84 |
| PC1 | −0.015 | 0.909 | 0.054 | 0.67 |
| PC2 | −0.107 | 0.403 | −0.056 | 0.66 |

***Figure S1. Swim-bout and freezing-bout rates (code output: Extra_bout_structure; data: trial_level_metrics)***

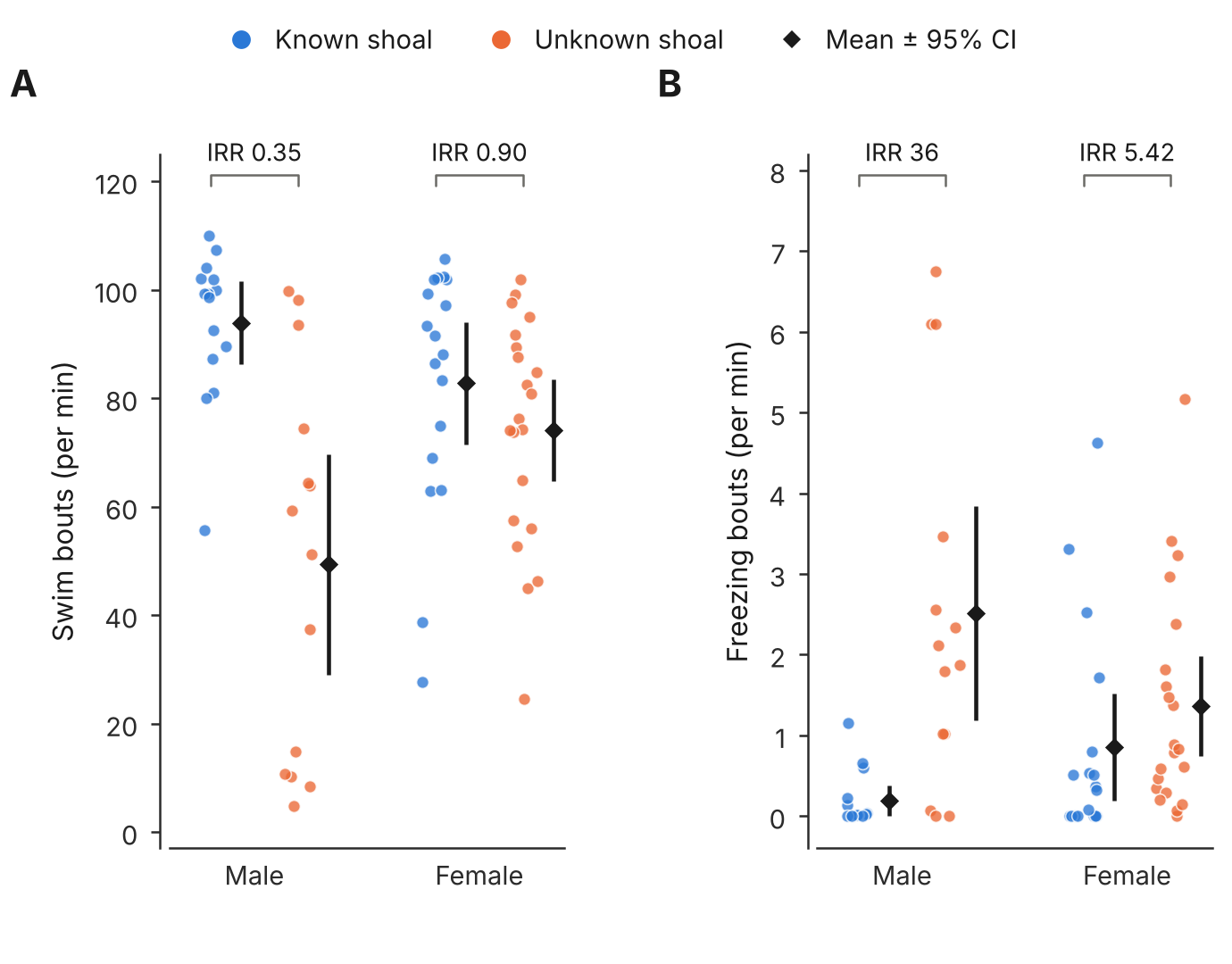

Swim bouts (A) and freezing bouts (B) per minute for each fish (points), by sex and treatment; diamonds and bars show mean ± 95% CI. Annotations give the incidence-rate ratio (Unknown vs Known) from the Poisson mixed model: the male values are the Treatment coefficients and the female values are derived from the Treatment and Treatment × Sex coefficients (Table S3).

***Figure S2. Principal component analysis of per-fish behavioural profiles***

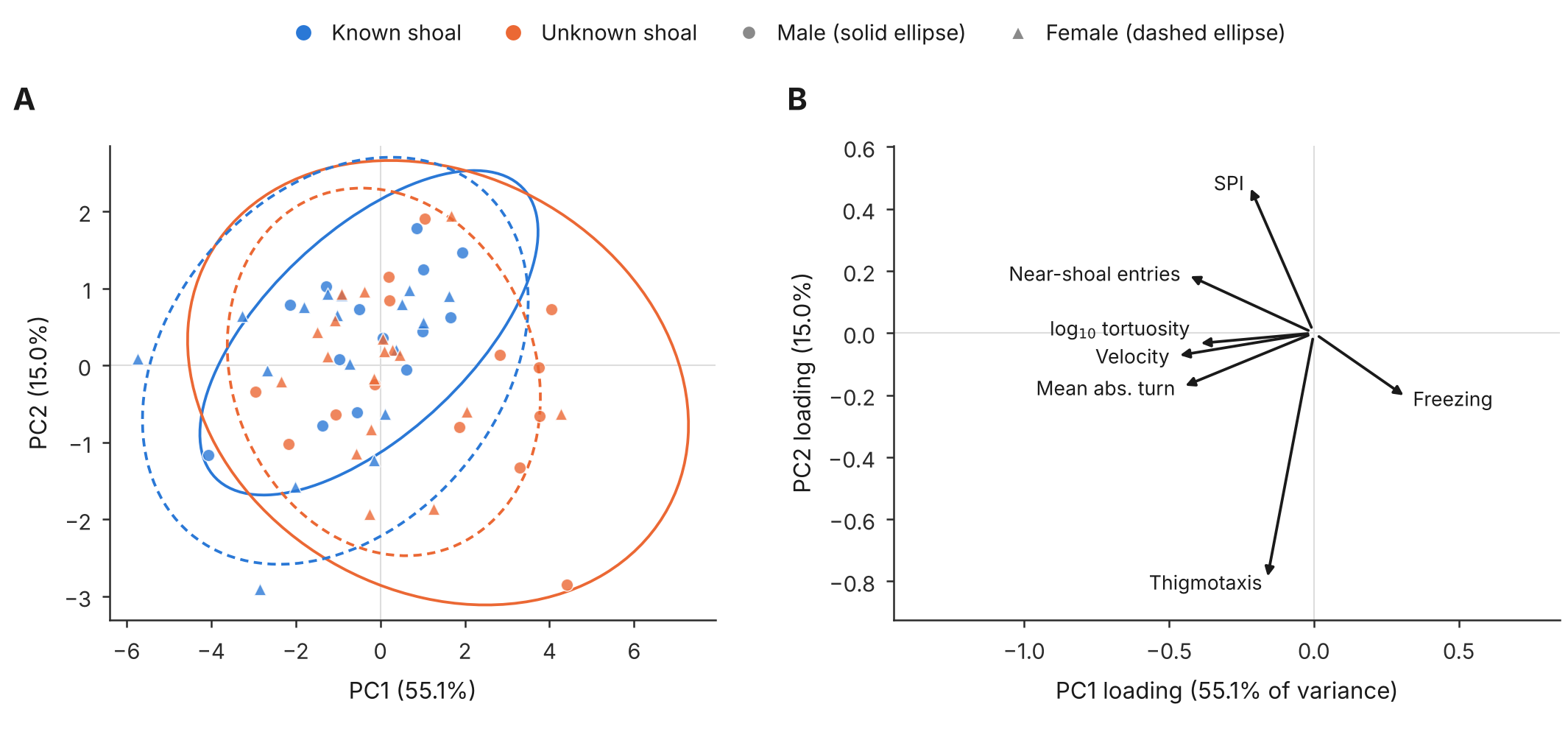

A) PC1 (55.1% of variance) against PC2 (15.0%) for individual fish with 95% ellipses per Treatment × Sex group (males, circles and solid ellipses; females, triangles and dashed ellipses). B) Loadings of the seven z-scored per-fish metrics (recomputed from per-fish means; matches the pipeline output). Freezing loads positively on PC1; velocity, mean absolute turn, near-shoal entries, tortuosity and SPI load negatively; thigmotaxis and, less so, SPI load mainly on PC2.

***Figure S3. Time courses of social preference and near-shoal entries***

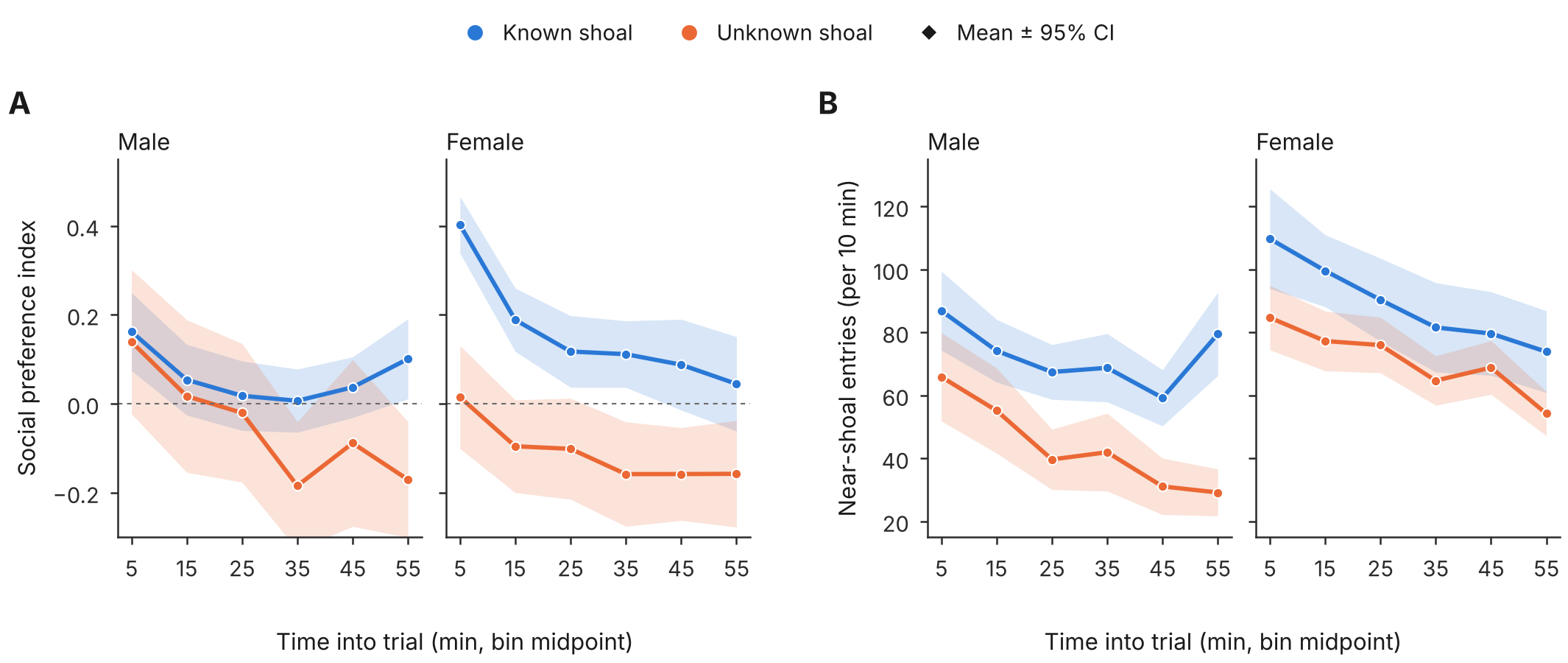

Social preference index (A) and entries into the near-shoal zone (B) in 10-min bins for males and females exposed to a Known (blue) or Unknown (orange) shoal; lines show mean ± s.e.m. of per-fish values.

***Table S9. Checks supporting each main conclusion***

| **Conclusion** | **Check** | **Result** |
| --- | --- | --- |
| Freezing increases in males | Fish as unit of replication; BH correction; random slopes; shoal random effect | Holds (P = 0.00090; BH 0.0045) (Tables S1, S5) |
| Same, independent of definition | Freezing threshold sensitivity | Significant at ≥ 0.3 cm s⁻¹, not at 0.2 cm s⁻¹; female effect never significant (Table S5) |
| Same, independent of threshold | Threshold-free HMM | Frozen-state occupancy 8% → 49% in males (Table S4) |
| Sex difference in behaviour | Treatment × Sex interactions | Not significant for freezing (P = 0.082), velocity, entries, SPI, thigmotaxis; significant for swim bouts and HMM straight-swim state; compositional check P = 0.045 (Tables S1, S3, S4) |
| Cortisol higher after Unknown shoal | Samples below the lowest standard treated as censored; weight, session, batch and shoal adjustment; exclusion of the two zero-standard fish | Censored regression (11 samples censored): overall 1.6-fold, P = 0.030; males 1.9-fold, P = 0.045. Only K18 and K19 censored: males 2.1-fold, P = 0.019. Extrapolated values: males 3.6-fold, P = 0.006; excluding K18 and K19: 1.85-fold, P = 0.078 (Tables S2, S5, S12) |
| Cortisol sex difference | Treatment × Sex interaction | Not significant (censored regression P = 0.40; extrapolated values P = 0.107); after a Known shoal, females tended to have higher cortisol than males (n.s.) (Tables S2, S12) |
| Cortisol and behaviour decoupled | Spearman and Pearson correlations | All \|ρ\| ≤ 0.13, P ≥ 0.32; within Treatment × Sex groups \|ρ\| ≤ 0.15, P ≥ 0.26 (Tables S8, S12) |
| Not a model-assumption artefact | glmmTMB refits with outcome-appropriate families | Male freezing effect significant in all four fits (P ≤ 0.0013); velocity and entries significant in males under Gamma and Poisson/negative binomial; SPI and thigmotaxis null; Treatment × Sex interaction significant only for velocity under the Gamma model (P = 0.039) (Table S10) |
| Not clustering by day, shoal or arena position | Random effects for recording day and stimulus shoal; day as fixed effect; arena position (quadrant) as fixed effect | Male freezing effect P = 0.0010–0.0022; with quadrant +11.9 points, P = 0.007; HMM state effects unchanged (Tables S11, S11b) |
| Power for interactions | Minimum detectable Treatment × Sex effect at 80% power | Freezing 15.6 percentage points; cortisol 0.76 log₁₀ (Table S15) |

***Table S10. glmmTMB refits of the behavioural models (code output: TableS10a, S10b, S10c)***

Same fixed effects as the primary models (Treatment × Sex + quadratic time) with random intercepts for session and fish. Estimates are on the link scale (identity for Gaussian; logit for binomial, beta-binomial and beta; log for Poisson, negative binomial and Gamma), so a negative log estimate is a ratio below 1. Male and female columns are Unknown − Known contrasts with Wald (asymptotic) P values; Gaussian glmmTMB P values are therefore slightly smaller than the Satterthwaite P values in Table S1. All 14 models converged with positive-definite Hessians. The plain binomial model is over-dispersed (Pearson dispersion 300) and its P values are anti-conservative; the beta-binomial and observation-level-random-effect versions account for this.

| **Outcome** | **Model** | **Males (est ± SE)** | **P** | **Females (est ± SE)** | **P** | **Treatment × Sex P** | **Converged** |
| --- | --- | --- | --- | --- | --- | --- | --- |
| Freezing | Gaussian (REML) | 14.5 ± 4.2 | 0.0005 | 4.78 ± 3.6 | 0.18 | 0.078 | yes |
| Freezing | Binomial (frames) | 4.72 ± 1.5 | 0.0013 | 3.48 ± 1.2 | 0.0046 | 0.52 | yes |
| Freezing | Binomial + OLRE | 8.39 ± 1.9 | 1.1e-05 | 4.54 ± 1.6 | 0.0058 | 0.12 | yes |
| Freezing | Beta-binomial | 3.15 ± 0.77 | 4.1e-05 | 1.25 ± 0.63 | 0.047 | 0.056 | yes |
| Entries near shoal | Gaussian (REML) | -28.8 ± 15 | 0.06 | -18.2 ± 13 | 0.17 | 0.6 | yes |
| Entries near shoal | Poisson + OLRE | -0.829 ± 0.3 | 0.0058 | -0.243 ± 0.26 | 0.35 | 0.14 | yes |
| Entries near shoal | Negative binomial (nbinom2) | -0.783 ± 0.29 | 0.0065 | -0.242 ± 0.25 | 0.33 | 0.15 | yes |
| SPI | Gaussian (REML) | -0.114 ± 0.16 | 0.47 | -0.268 ± 0.14 | 0.049 | 0.46 | yes |
| SPI | Beta (SPI rescaled to 0-1) | -0.453 ± 0.41 | 0.27 | -0.727 ± 0.35 | 0.039 | 0.61 | yes |
| Thigmotaxis | Gaussian (REML) | 6.22 ± 5.5 | 0.26 | -5.64 ± 4.8 | 0.24 | 0.11 | yes |
| Thigmotaxis | Binomial + OLRE | 0.259 ± 0.28 | 0.35 | -0.247 ± 0.24 | 0.31 | 0.17 | yes |
| Thigmotaxis | Beta-binomial | 0.231 ± 0.24 | 0.33 | -0.209 ± 0.21 | 0.31 | 0.16 | yes |
| Velocity | Gaussian (REML) | -1.48 ± 0.68 | 0.029 | -0.543 ± 0.58 | 0.35 | 0.3 | yes |
| Velocity | Gamma (log link) | -0.583 ± 0.19 | 0.0021 | -0.0659 ± 0.16 | 0.69 | 0.039 | yes |

***Table S11. Additional checks on the behavioural models (code output: TableS11_extra_checks)***

Males: Unknown − Known estimate and P from linear mixed models with fish random intercept plus the stated terms. Residual shape refers to the Gaussian model with a fish random intercept.

| **Outcome** | **Check** | **Specification** | **Male estimate** | **P** | **Note** |
| --- | --- | --- | --- | --- | --- |
| Freezing | Residual shape | (1\|fish) | — | — | skew=0.93 excess kurtosis=2.34 |
| Freezing | Cluster RE | (1\|fish_id)+(1\|Day) | 14.5 | 0.001 | P(TxS)=0.082 |
| Freezing | Cluster RE | (1\|fish_id)+(1\|Day)+(1\|Shoal) | 14.5 | 0.0022 | P(TxS)=0.082 |
| Freezing | Day fixed effect | + Day (fixed) + (1\|fish_id) | 14.9 | 0.0016 | P(TxS)=0.079 |
| Entries near shoal | Residual shape | (1\|fish) | — | — | skew=0.23 excess kurtosis=1.51 |
| Entries near shoal | Cluster RE | (1\|fish_id)+(1\|Day) | -29 | 0.066 | P(TxS)=0.600 |
| Entries near shoal | Cluster RE | (1\|fish_id)+(1\|Day)+(1\|Shoal) | -29 | 0.086 | P(TxS)=0.600 |
| Entries near shoal | Day fixed effect | + Day (fixed) + (1\|fish_id) | -32 | 0.054 | P(TxS)=0.580 |
| SPI | Residual shape | (1\|fish) | — | — | skew=-0.10 excess kurtosis=1.13 |
| SPI | Cluster RE | (1\|fish_id)+(1\|Day) | -0.114 | 0.47 | P(TxS)=0.462 |
| SPI | Cluster RE | (1\|fish_id)+(1\|Day)+(1\|Shoal) | -0.105 | 0.54 | P(TxS)=0.477 |
| SPI | Day fixed effect | + Day (fixed) + (1\|fish_id) | -0.171 | 0.31 | P(TxS)=0.564 |
| Thigmotaxis | Residual shape | (1\|fish) | — | — | skew=0.56 excess kurtosis=1.43 |
| Thigmotaxis | Cluster RE | (1\|fish_id)+(1\|Day) | 5.53 | 0.33 | P(TxS)=0.127 |
| Thigmotaxis | Cluster RE | (1\|fish_id)+(1\|Day)+(1\|Shoal) | 5.53 | 0.36 | P(TxS)=0.127 |
| Thigmotaxis | Day fixed effect | + Day (fixed) + (1\|fish_id) | 3.03 | 0.6 | P(TxS)=0.222 |
| Velocity | Residual shape | (1\|fish) | — | — | skew=-0.52 excess kurtosis=9.93 |
| Velocity | Cluster RE | (1\|fish_id)+(1\|Day) | -1.48 | 0.035 | P(TxS)=0.301 |
| Velocity | Cluster RE | (1\|fish_id)+(1\|Day)+(1\|Shoal) | -1.48 | 0.049 | P(TxS)=0.301 |
| Velocity | Day fixed effect | + Day (fixed) + (1\|fish_id) | -1.51 | 0.042 | P(TxS)=0.304 |
| Velocity | Log-scale LMM | log(velocity_mean) | -0.627 | 0.0025 | P(TxS)=0.041; resid skew=-2.29 kurt=25.35 |
| Entries near shoal | Log-scale LMM | log1p(entries_near_shoal) | -0.913 | 0.0062 | P(TxS)=0.135; resid skew=-1.43 kurt=11.54 |

***Table S11b. Arena position (code output: TableS11b_quadrant_sensitivity)***

Quadrant is unbalanced across Treatment × Sex (for example, no Unknown-shoal male was tested in the right-top arena), so it was added as a fixed effect. Behaviour: linear mixed models as in Table S1 with (1|Session) + (1|fish_id); HMM: beta regression of occupancy (logit scale).

| **Outcome** | **Model** | **Male estimate** | **P** | **Female estimate** | **P** | **Treatment × Sex P** | **Quadrant P** |
| --- | --- | --- | --- | --- | --- | --- | --- |
| Freezing | primary | 14.5 | 0.0012 | 4.78 | 0.2 | 0.082 | — |
| Freezing | + quadrant (fixed) | 11.9 | 0.0068 | 3.54 | 0.32 | 0.12 | 0.03 |
| Velocity | primary | -1.48 | 0.037 | -0.543 | 0.37 | 0.3 | — |
| Velocity | + quadrant (fixed) | -1.59 | 0.031 | -0.499 | 0.41 | 0.23 | 0.27 |
| Entries near shoal | primary | -28.8 | 0.072 | -18.2 | 0.18 | 0.6 | — |
| Entries near shoal | + quadrant (fixed) | -29.9 | 0.064 | -14.6 | 0.28 | 0.44 | 0.048 |
| SPI | primary | -0.114 | 0.48 | -0.268 | 0.058 | 0.46 | — |
| SPI | + quadrant (fixed) | -0.0758 | 0.59 | -0.187 | 0.12 | 0.53 | 2.5e-05 |
| Thigmotaxis | primary | 6.22 | 0.28 | -5.64 | 0.25 | 0.11 | — |
| Thigmotaxis | + quadrant (fixed) | 4.87 | 0.42 | -5.24 | 0.3 | 0.18 | 0.59 |
| HMM frozen (logit) | primary | 1.75 | 9e-05 | 0.649 | 0.072 | 0.059 | — |
| HMM frozen (logit) | + quadrant (fixed) | 1.82 | 8.8e-05 | 0.681 | 0.065 | 0.056 | — |
| HMM straight (logit) | primary | -1.47 | 3.3e-08 | -0.111 | 0.61 | 7.7e-05 | — |
| HMM straight (logit) | + quadrant (fixed) | -1.34 | 2e-07 | -0.095 | 0.65 | 0.00015 | — |

***Table S12. Cortisol sensitivity analyses and within-group cortisol–behaviour associations (code output: TableS12a, S12b, S12c)***

Fish with the two lowest cortisol values (K19, 5.3 pg g⁻¹; K18, 18.9 pg g⁻¹; both Known-shoal males; next lowest 194.9 pg g⁻¹) were excluded in turn. This table uses the extrapolated values. Overall effect = Unknown − Known averaged over sexes (Type-III F test); fold change from log₁₀ differences. The primary analysis in the main text treats all 11 samples below the lowest standard as left-censored (last row of the second table below; Table S2b).

| **Excluded** | **n** | **Overall fold** | **P** | **Male fold [95% CI]** | **P** | **Female P** | **Treatment × Sex P** |
| --- | --- | --- | --- | --- | --- | --- | --- |
| none (extrapolated values) | 65 | 2.17 | 0.014 | 3.57 [1.46, 8.74] | 0.006 | 0.51 | 0.11 |
| K19 | 64 | 1.82 | 0.026 | 2.50 [1.15, 5.42] | 0.021 | 0.43 | 0.23 |
| K18+K19 | 63 | 1.56 | 0.056 | 1.85 [0.93, 3.66] | 0.078 | 0.36 | 0.47 |

Other tests of the male contrast with all fish: HC3 robust P = 0.040; permutation P = 0.032; Wilcoxon P = 0.071 (females P = 0.30). Residuals of the linear model on extrapolated values are left-skewed (Shapiro–Wilk P < 0.001), driven by these two fish; maximum Cook's distance 0.26.

Treating non-detects as censored. K18 and K19 read at the zero-standard level in the ELISA (B/B0 ≈ 100%), below the lowest standard (50 pg mL⁻¹, i.e. 250 pg mL⁻¹ after the 1:5 dilution), so their concentrations are extrapolated. They were set to the detection limit (substitution) or treated as left-censored in a Tobit regression. The last scenario applies the same rule to all 11 samples measured below the lowest standard. That last Tobit row is the primary analysis.

| **Scenario** | **Censored** | **Method** | **Male fold [95% CI]** | **Male P** | **Female P** | **Overall P** | **Treatment × Sex P** |
| --- | --- | --- | --- | --- | --- | --- | --- |
| Extrapolated values kept (no censoring) | 0 | substitution + linear model | 3.57 [1.46, 8.74] | 0.006 | 0.51 | 0.014 | 0.11 |
| K18, K19 set to detection limit | 2 | substitution + linear model | 1.97 [1.03, 3.77] | 0.041 | 0.36 | 0.035 | 0.37 |
| K18, K19 set to detection limit | 2 | left-censored (Tobit) regression | 2.13 [1.13, 4.01] | 0.019 | 0.35 | 0.018 | 0.27 |
| All samples below lowest standard set to detection limit | 11 | substitution + linear model | 1.51 [0.87, 2.63] | 0.14 | 0.23 | 0.060 | 0.79 |
| All samples below lowest standard set to detection limit (primary) | 11 | left-censored (Tobit) regression | 1.92 [1.01, 3.63] | 0.045 | 0.31 | 0.030 | 0.40 |

| **Metric** | **n** | **Pooled ρ** | **P** | **ρ within Treatment × Sex** | **P** | **ρ (cortisol also weight-adjusted)** | **P** |
| --- | --- | --- | --- | --- | --- | --- | --- |
| freezing_pct | 63 | 0.126 | 0.32 | -0.0182 | 0.89 | -0.00579 | 0.96 |
| SPI | 63 | -0.107 | 0.41 | -0.089 | 0.49 | -0.0955 | 0.46 |
| entries_near_shoal | 63 | -0.00197 | 0.99 | 0.107 | 0.4 | 0.111 | 0.39 |
| velocity_mean | 63 | 0.0307 | 0.81 | 0.145 | 0.26 | 0.152 | 0.23 |
| thigmotaxis_pct | 63 | 0.0501 | 0.7 | -0.0023 | 0.99 | 0.00739 | 0.95 |
| PC1 | 63 | -0.0147 | 0.91 | -0.1 | 0.43 | -0.102 | 0.42 |
| PC2 | 63 | -0.107 | 0.4 | 0.00878 | 0.95 | -0.00322 | 0.98 |

***Table S13. HMM occupancy: sensitivity of the beta regression (code output: TableS13_hmm_beta_sensitivity)***

Occupancy proportions were clamped to [10⁻⁴, 1 − 10⁻⁴] in the primary analysis (two fish had exactly zero frozen-state occupancy and 13 had < 1%). The alternative squeezes proportions as (p(n − 1) + 0.5)/n. Logit estimates are Unknown − Known in males.

| **State** | **Analysis** | **Male logit** | **Male P** | **Female P** | **Treatment × Sex P** |
| --- | --- | --- | --- | --- | --- |
| frozen | clamp 1e-4 (primary) | 1.75 | 9e-05 | 0.072 | 0.059 |
| frozen | Smithson-Verkuilen | 1.64 | 0.0001 | 0.15 | 0.033 |
| frozen | Wilcoxon per sex | — | 0.0016 | 0.1 | — |
| straight | clamp 1e-4 (primary) | -1.47 | 3.3e-08 | 0.61 | 7.7e-05 |
| straight | Smithson-Verkuilen | -1.44 | 3.7e-08 | 0.61 | 8.4e-05 |
| straight | Wilcoxon per sex | — | 0.00038 | 0.56 | — |
| turning | clamp 1e-4 (primary) | -0.697 | 0.00042 | 0.31 | 0.035 |
| turning | Smithson-Verkuilen | -0.66 | 0.00052 | 0.3 | 0.041 |
| turning | Wilcoxon per sex | — | 0.058 | 0.39 | — |

***Table S14. Stimulus shoal and recording day by treatment (fish tested, n = 70; shoal coded case-insensitively; code output: TableS14_shoal_day_by_treatment)***

| **Stimulus shoal** | **Known** | **Unknown** |
| --- | --- | --- |
| stim1 | 4 | 6 |
| stim2 | 2 | 8 |
| stim3 | 4 | 2 |
| stim4 | 6 | 14 |
| stim5 | 12 | 2 |
| stim6 | 6 | 4 |

| **Recording day** | **Known** | **Unknown** |
| --- | --- | --- |
| 20240920 | 4 | 4 |
| 20240926 | 8 | 8 |
| 20240927 | 4 | 8 |
| 20241003 | 4 | 8 |
| 20241004 | 6 | 0 |
| 20241017 | 8 | 8 |

Shoals and days are unevenly distributed across treatments (for example stim5: 12 Known, 2 Unknown; stim4: 6 Known, 14 Unknown; 4 October 2024: Known only), so shoal and day were examined as random effects (Tables S5, S11).

***Table S15. Minimum detectable Treatment × Sex interaction (code output: TableS15_interaction_MDE)***

Post hoc sensitivity, not observed power: (t₀.₉₇₅ + t₀.₈₀) × SE of the interaction coefficient, i.e. the smallest interaction detectable with 80% power at α = 0.05 given the observed standard errors.

| **Outcome** | **Observed interaction** | **SE** | **Minimum detectable (80% power)** |
| --- | --- | --- | --- |
| freezing_pct | -9.68 | 5.49 | 15.6 |
| velocity_mean | 0.933 | 0.894 | 2.54 |
| entries_near_shoal | 10.7 | 20.3 | 57.7 |
| SPI | -0.154 | 0.208 | 0.591 |
| thigmotaxis_pct | -11.9 | 7.32 | 20.8 |
| log10 cortisol/g | -0.434 | 0.265 | 0.756 |

***Table S16. Thigmotaxis with and without the wall facing the stimulus shoal and with frozen frames removed (code output: TableS16_thigmotaxis_sensitivity)***

Mean percentage of time within 1.5 cm of a wall, and Unknown − Known contrasts (percentage points) from the same mixed model as Table S1. Non-frozen variants use only tracked frames in which the fish was not freezing, so that a fish freezing beside a wall is not counted as wall-seeking.

| **Measure** | **Mean %** | **Males** | **P** | **Females** | **P** | **Treatment × Sex P** |
| --- | --- | --- | --- | --- | --- | --- |
| Any wall (primary) | 43.0 | +6.2 | 0.28 | -5.6 | 0.25 | 0.11 |
| Excluding shoal-facing wall | 30.4 | +6.1 | 0.27 | +0.12 | 0.98 | 0.4 |
| Non-frozen frames only | 43.4 | +6.4 | 0.26 | -4.3 | 0.38 | 0.14 |
| Non-frozen frames, excluding shoal-facing wall | 30.7 | +6.3 | 0.24 | +0.67 | 0.88 | 0.41 |

***Table S17. Swim-bout rate at different peak-velocity cut-offs (code output: TableS17_swim_bout_cutoff_sensitivity)***

Poisson mixed model with observation-level random effect and observation-time offset; bouts are peaks of smoothed velocity above the cut-off (primary 1.5 cm s⁻¹). IRR = Unknown / Known in males; interaction IRR is the additional ratio in females.

| **Cut-off (cm s⁻¹)** | **Mean bouts per fish** | **Males IRR [95% CI]** | **P** | **Treatment × Sex IRR** | **P** |
| --- | --- | --- | --- | --- | --- |
| 1.0 | 4420 | 0.39 [0.27, 0.56] | 2.5e-07 | 2.34 | 0.00046 |
| 1.5 | 4260 | 0.35 [0.24, 0.52] | 2.4e-07 | 2.55 | 0.00044 |
| 2.0 | 4070 | 0.36 [0.24, 0.54] | 8.8e-07 | 2.51 | 0.00091 |
